# Omics profiling identifies MAPK/ERK pathway as a gatekeeper of nephron progenitor metabolism

**DOI:** 10.1101/2021.09.27.461969

**Authors:** Hyuk Nam Kwon, Kristen Kurtzeborn, Xing Jin, Abigail Loh, Nathalie Escande-Beillard, Bruno Reversade, Sunghyouk Park, Satu Kuure

## Abstract

Nephron endowment is defined by fetal kidney growth and critically dictates renal health in adults. Despite the advances in understanding the molecular regulation of nephron progenitors, the causes for low congenital nephron count and contribution of basic metabolism to nephron progenitor biology remain poorly understood. Here we characterized the metabolic consequences of MAPK/ERK-deficiency in nephron progenitors, whose maintenance and propagation in developing kidney critically depends on ERK activation. Our LC/MS-based metabolomics profiling identified 42 reduced metabolites, of which 26 were further supported by *in vivo* transcriptional characterization of MAPK/ERK-deficient nephron progenitors. This revealed a severe shortage of energy and nucleotide biosynthesis precursors, blockage in glycolysis and diminished pyruvate and proline metabolism. Utilization of *in vitro* kidney cultures demonstrated a dosage-specific function for glycolytic pyruvate as an energy source that controls the shape of the ureteric bud tip kwon to serve as a niche for nephron progenitor regulation. Analysis of the proline biosynthesis effects in developing kidney *in vivo* revealed premature loss of nephron progenitor maintenance in the absence of *Pycr1*/2 functions. Our results suggest that MAPK/ERK-dependent nephron progenitor metabolism functionally contributes to progenitor preservation by controlling pyruvate availability and proline metabolism in developing kidneys.

## INTRODUCTION

The differentiation capacity of embryonic nephron progenitors (NP) is a crucial factor for renal health later in life as the extent of fetal nephrogenesis determines an individual’s final nephron number, and low nephron count predicts a higher risk for renal complications (1, 2). Embryonic kidney development is orchestrated by inter- and intracellular signaling taking place between two mesodermal derivatives, the ureteric bud (UB) and metanephric mesenchyme (MM) (3, 4). The process of UB branching patterns and expands kidney in size while the MM hosts NPs, which give rise to all segments of functional nephron (5). Together, the UB and MM form the NP niche, where balanced propagation and differentiation of MM-residing progenitors are guided by niche-intrinsic and extrinsic cues (6).

The total NP lifespan is limited to the fetal period in humans and to the first postnatal days of life in mouse (7–9). The molecular regulation of NP maintenance and differentiation has been extensively studied (10, 11), but the factors contributing to the cessation of the nephrogenic program are only evolving and suggest involvement of changes in signal transduction activities (12–14). Moreover, information is only emerging about metabolic events and especially mitochondrial metabolism, underlying the cell type-specific behaviors of NPs during their propagation and differentiation switches (15, 16). A recent study demonstrated higher glycolysis dependance in younger versus older NPs and showed that glycolysis inhibition promotes differentiation being in line with the requirement for aerobic glycolysis (also known as Warburg effect) in maintenance of pluripotency (17, 18). Additionally, Von Hippel-Lindau (VHL) tumor suppressor appears important for mitochondrial respiration in NPs, which show mild differentiation deficiency in the NP-specific absence of *Vhl* (19).

Although one of the major energy precursors for mitochondria, namely pyruvate, is essential for normal early embryonic development and is suggested to contribute to the cell fate switches in embryonic stem cells (20–22), general understanding of its functions in organogenesis is missing. The last step of pyruvate production is regulated by the embryonic kidney expressed glycolytic enzyme pyruvate kinase isozyme M2 (PKM2), whose function is controlled by the mitogen-activated protein kinase/extracellular signal-regulated kinase (MAPK/ERK) and whose reduced activity is shown to protect against kidney injury (23, 24). Similar to MAPK/ERK, PKM2 regulates cell cycle in embryonic cardiomyocytes (25–28).

We have recently demonstrated the importance of MAPK/ERK activity to the transmission of extracellular signals for NP regulation, where it on one hand maintains NP identity and on the other hand propels differentiation in nephron precursors (4, 11, 29). Interestingly, MAPK/ERK target p53 contributes to metabolic fitness in propagating NPs (30). The p53 target Rrm2b, encoding a catalytic subunit of ribonucleotide reductase (p53R2), is required for nephron function and interestingly interacts with mitochondrial proline synthesis enzymes pyrroline-5-carboxylate reductase (PYCR) 1 and 2 (31–33). Proline metabolism is heavily utilized in cancer cells and deficiencies in its metabolism are causative for many congenital disorders including different types of *cutis laxa* with rare renal manifestation (34–36). Emerging evidence also supports a regulatory role for proline in the maintenance of pluripotency at least in embryonic stem cells (37, 38). Single and double knockout studies of *Pycr1* and -*2* demonstrated essential involvement of proline metabolism in neuronal differentiation (39, 40), but its other organ- and tissue-specific requirements remain elusive.

We have utilized a systems biology approach to study the cellular functions that MAPK/ERK activity effectuates in NPs. Our combined RNA sequencing and metabolomics characterization of control and MAPK/ERK-deficient NPs revealed cellular metabolism as one of the most important contributing factors in their regulation. These results establish that the MAPK/ERK pathway guides nephrogenesis in developing kidney through regulating energy metabolism in NPs.

## RESULTS

### Transcriptional profiling identifies mitochondrial metabolism as the most MAPK/ERK-dependent cellular process in nephron progenitors

Recently, our group utilized a genetic approach to reveal critical roles for NP-specific MAPK/ERK signaling in progenitor maintenance and propagation (20). NP-specific loss of MAPK/ERK activation due to conditional inactivation of both *Mek* genes (Six2-TGC^tg/+^;*Mek1*^fl/fl^;*Mek2*^-/-^) results in rapid NP deprivation due to cell-intrinsic proliferation defects and molecular changes in the niche composition. Transcriptional profiling of MAPK/ERK-deficient NPs isolated from E13.5 kidneys (before the onset of significant NP loss) identified over 5,000 differentially expressed genes (DEGs), which represent transcripts with significant effects on mitochondrial energy metabolism (Kurtzeborn et al, manuscript in revision). To further investigate the mitochondrial metabolism in NP populations, we now performed the biological interpretation analysis of the NP RNA-Seq data using the Cytoscape plug-in ClueGO (41, 42).

An overview of the gene ontology (GO) analysis of DEGs (Padj<0.05 and │log2fold change│≥2) identified in MAPK/ERK-deficient NPs is shown in Figure 1A. This shows that oxidative phosphorylation (18.87%) and mitochondrion (11.32%), which represent mitochondrial metabolism, account for a total of 30.19% of cellular functions affected by MAPK/ERK-deficiency and demonstrates that mitochondrial metabolism represents the largest portion (nearly one-third) of cellular pathways that depend on MAPK/ERK. Further characterization of down-regulated DEG results by integrating “cellular component” and Kyoto Encyclopedia of Genes and Genomes (KEGG) database pathway (43, 44) demonstrated that mitochondrial inner membrane (25.0%) and mitochondrion (15.0%) account 40.0% of the down-regulated transcripts (Figure 1B) supporting diminished mitochondrial metabolism in the absence of MAPK/ERK activation. The up-regulated genes were so scattered and few that their pathway analysis failed to provide a meaningful outcome (Table S1).

**Figure 1.**
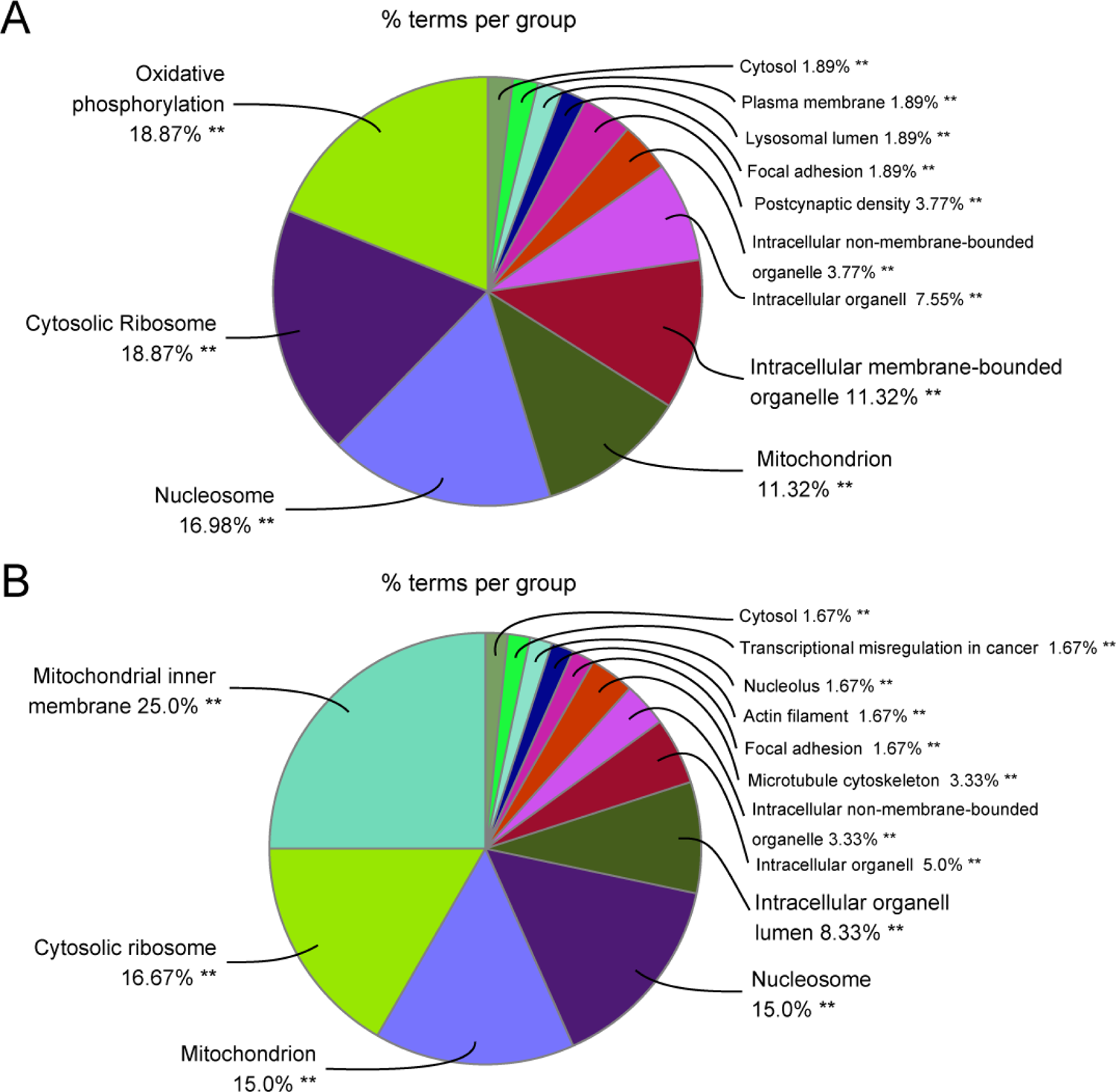
ClueGO analysis of differentially expressed transcripts in MAPK/ERK-deficient nephron progenitors. Overview pie chart of cellular component and KEGG pathway analyses of the (**A**) total and (**B**) down-regulated differentially expressed genes (DEGs) between control and MAPK/ERK-deficient NPs. The percentage of genes per term is shown in each group. GO analysis was performed by Cytoscape plug-in application ClueGO. The pie chart shows the enriched signaling pathway categories based on the kappa coefficient of 0.5.

**Table 1.**
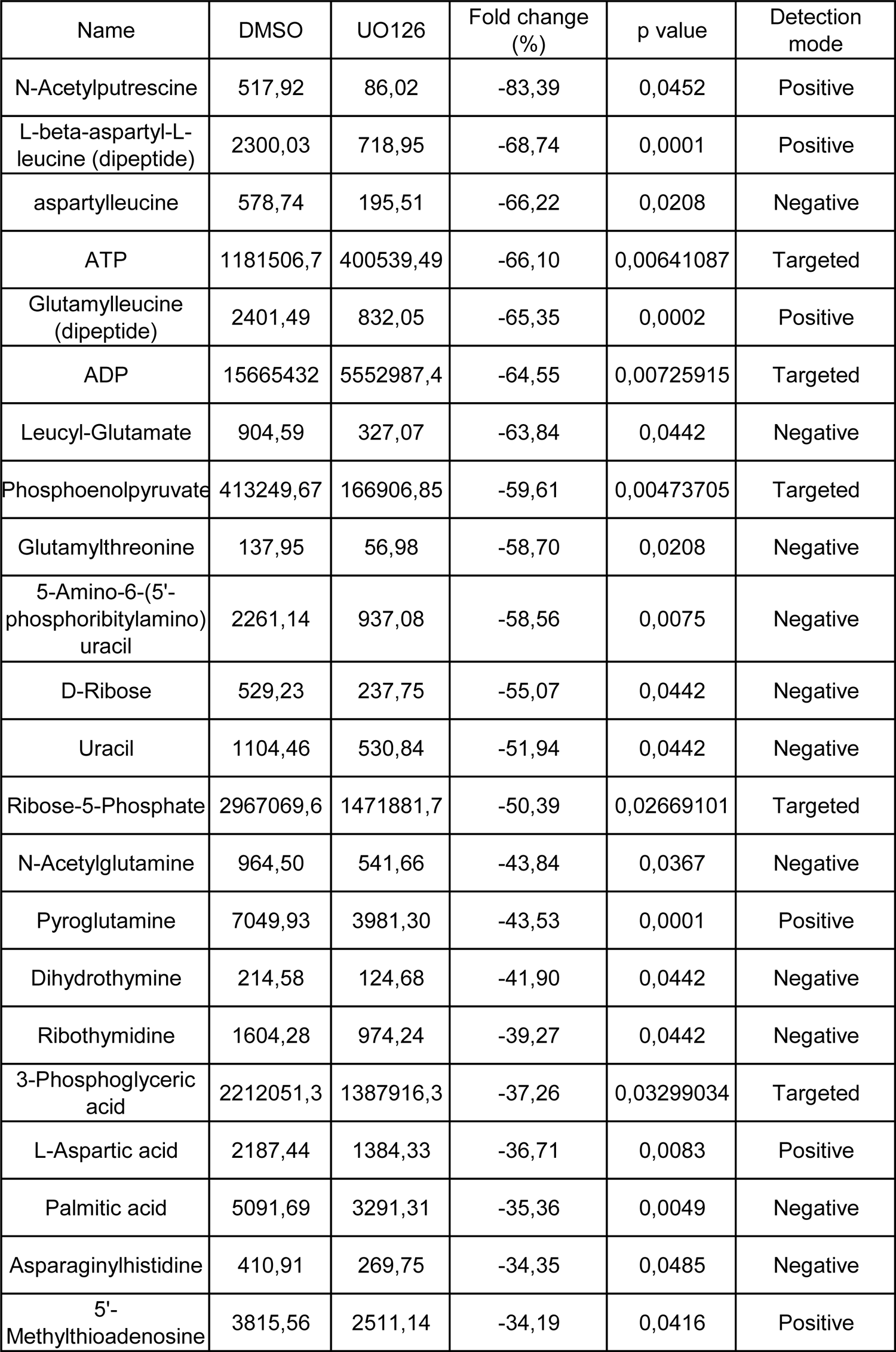
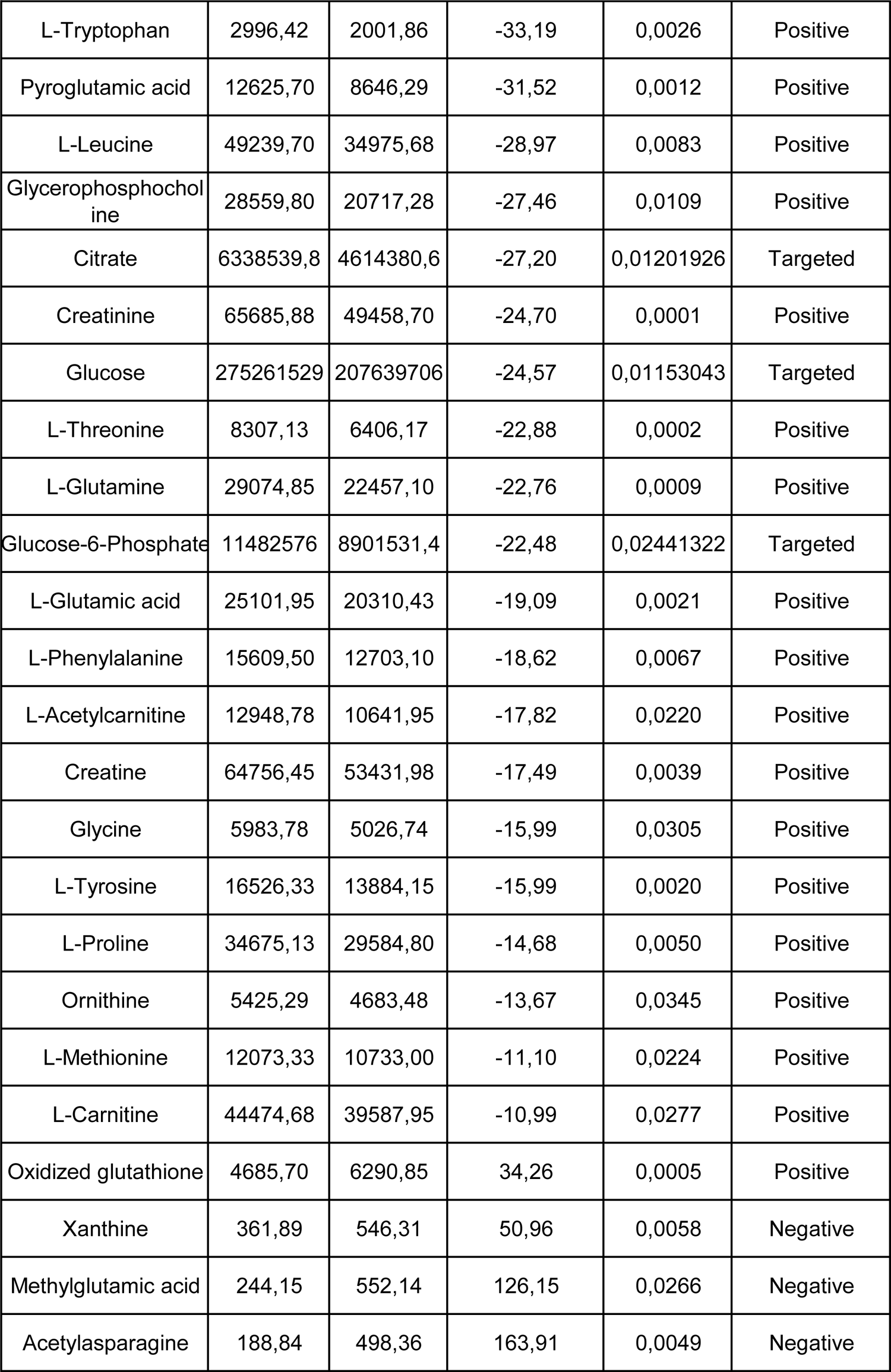
Metabolite marker identification

### Metabolomics analysis of MAPK/ERK-deficiency

We next applied the LC/MS-based metabolomics approach to assess MAPK/ERK-affected metabolism in more detail. An untargeted metabolomics approach was applied to the embryonic kidney mesenchyme cell line (mK4) treated with the MEK inhibitor U0126 to mimic genetic MAPK/ERK-deficiency *in vitro*. The LC/MS data of the non-treated cells, DMSO-treated cells, and the MEK-inhibited cells were analyzed by the partial least square’s discriminant analysis (PLS-DA), a multivariate statistical analysis technique. As a result, each group showed clear discrimination in both positive and negative detection mode, while clustering well within their treatment group (Figure 2). Comparison of the untargeted MS2 spectrum patterns with an online MS database revealed a total of 38 significantly different metabolites, summarized in Table 1. Of these, the majority of metabolites were down-regulated and only four (acetylasparagine, methylglutamic acid, xanthine, and oxidized glutathione) were up-regulated.

**Figure 2.**
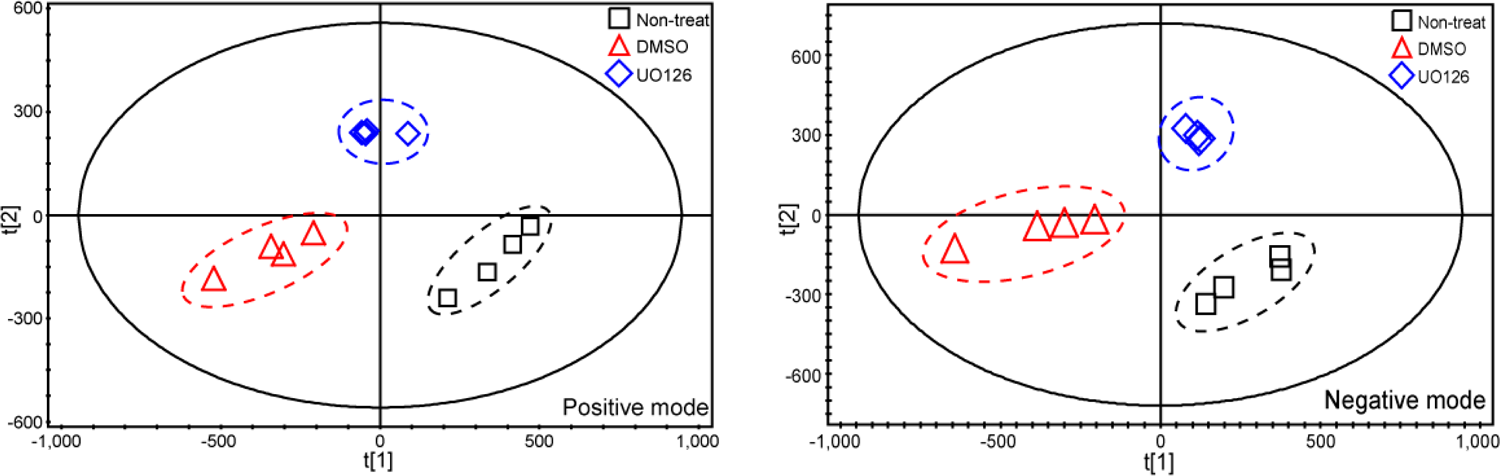
Partial least squares discriminant analysis (PLS-DA) on positive and negative mode metabolites identified in mK4 cell upon MEK inhibition by LC/MS. PLS-DA score plot for m/z values of (**A**) positive (R²X = 0.92; R²Y = 0.991; Q²Y = 0.968; n = 1 + 4 components) and (**B**) negative (R²X = 0.817; R²Y = 0.999; Q²Y = 0.992; n = 1 + 1 components) detection mode of LC/MS data. Each symbol represents the metabolic profile of an individual sample on each group (n=4). Black box indicates the non-treated group, red triangle indicates the DMSO treated group, and blue rhombus indicates the MEK inhibitor UO126 treated group. In both detection modes, MEK-inhibited cells, where MAPK/ERK activation is abolished share the most similar metabolic profile.

We next performed targeted metabolomics analysis aiming at identification of metabolic intermediates from glycolysis and mitochondrial metabolism, and quantification of principally altered metabolites. This revealed eight statistically different intermediate metabolites upon MAPK/ERK-deficiency: ATP, ADP, phosphoenolpyruvate, ribose-5-Phosphate, 3-phosphoglyceric acid, citrate, glucose, and glucose-6-phosphate, which all are down-regulated (Table 1). This quantitative intermediate metabolite result is well in line with our RNA-Seq results (Kurtzeborn et al, manuscript in revision) further analyzed here (Fig. 1), which indicated down-regulation of mitochondrial functions.

Metabolic set enrichment analysis (MSEA) of all identified metabolites by MetaboAnalyst revealed pentose phosphate pathway, Warburg effect, Glycine/Serine metabolism, and Purine metabolism as the most altered metabolic pathways in the absence of MAPK/ERK activation (Figure 3A). Visualization of the actual metabolite level changes by heatmaps shows that most of the metabolic pathways, but particularly energy-expensive metabolisms including nucleotide synthesis, aerobic glycolysis, and Glycine/Serine metabolism, were down-regulated upon MAPK/ERK-deficiency (Figure 3B). Table 2 summarizes MSEA analysis results and includes a list of the contributing metabolites.

**Figure 3.**
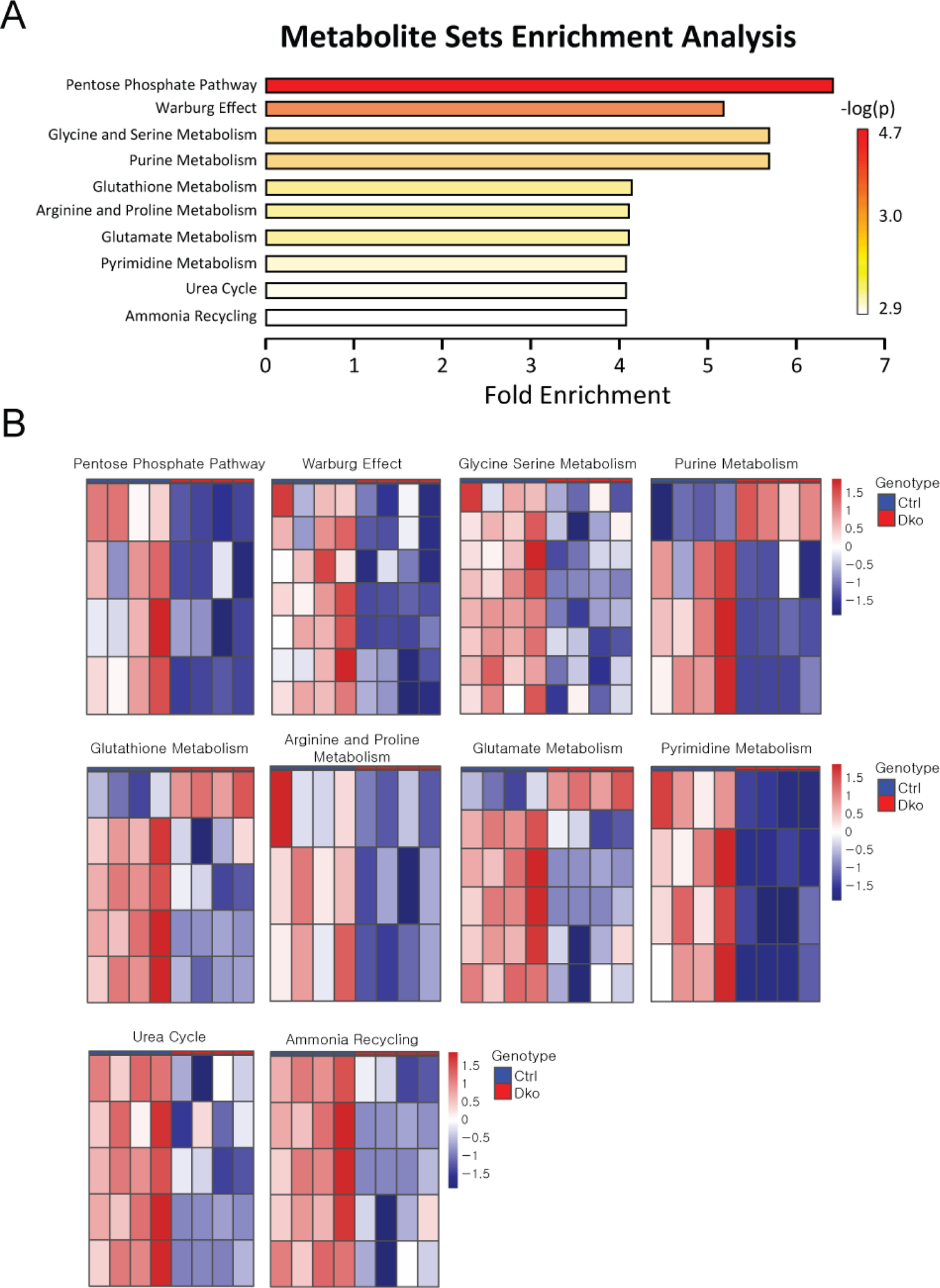
Metabolic Set Enrichment Analysis (MSEA) and their heatmap analyses. **(A)** All 46 MAPK/ERK-dependent metabolites identified by LC/MS were applied to MSEA to distinguish the most affected metabolic pathways. The horizontal bar plots are ranked by p value. The color gradient is an indicator of how strongly a metabolic pathway changes, with the strongest changes being red and lower changes white. (**B**) The heatmap analysis of the mRNA changes contributing to the differential pathways identified by MSEA.

**Table 2.** Metabolic set enrichment analysis (MSEA) of identified metabolites

| **<u>Metabolic changes</u>** | **<u>Total Comp</u>** | **<u>Hits</u>** | **<u>Raw p</u>** | **<u>Contributing metabolites</u>** |
| --- | --- | --- | --- | --- |
| Glycine and Serine Metabolism | 59 | 8 | 0,0007233 | 3PG, ADP, Glycine, Methionine, Glutamate, Threonine, ADP, Ornithinine |
| Warburg Effect | 58 | 7 | 0,0001962 | 3PG, R5P, G6P, Citrate, ADP, Glutamate, Glutamine |
| Purine Metabolism | 74 | 7 | 0,0007233 | Glutamine, ADP, R5P, Xanthine |
| Arginine and Proline Metabolism | 53 | 6 | 0,0010469 | Proline, Creatine, N-Acetyl putrescine |
| Glutamate Metabolism | 49 | 6 | 0,0010469 | Aspartate, Glycine, ADP, Glutamine, Glutamate, Oxidized Glutathione |
| Glutathione Metabolism | 21 | 5 | 0,0010464 | Pyroglutamic acid, ADP, Glutamate, Glycine, Oxidized Glutathione |
| Urea Cycle | 29 | 5 | 0,0010482 | Glutamine, Glutamate, ADP, Ornithine, Aspartate |
| Ammonia Recycling | 32 | 5 | 0,0010486 | Glutamine, Glutamate, ADP, Aspartate, Glycine |
| Pentose Phosphate Pathway | 29 | 4 | 2,31E-05 | D-ribose, R5P, G6P, ADP |
| Pyrimidine Metabolism | 59 | 4 | 0,0010478 | Glutamine, ADP, Uracil, Dihydrothymine |

### Integration of metabolomics and RNA-Seq results to the systems biology approach

To conduct a more comprehensive analysis at the level of whole systems biology, we next attempted to integrate the information from our metabolomics results and recent RNA-Seq data. The basic idea of metabolic pathways is a kind of catabolism regulated by enzymes, where these metabolic pathways are composed of regulating enzymes and the metabolites as end products. Here, we have obtained quantitative metabolite level changes in MAPK/ERK-deficient cells through a metabolomics approach and gene expression level changes in MAPK/ERK-deficient nephron progenitors by RNA-Seq. The mutually meaningful relationship between metabolite and gene expression levels was investigated by integrating the two analysis results, and the combined results are summarized in Figure 4.

**Figure 4.**
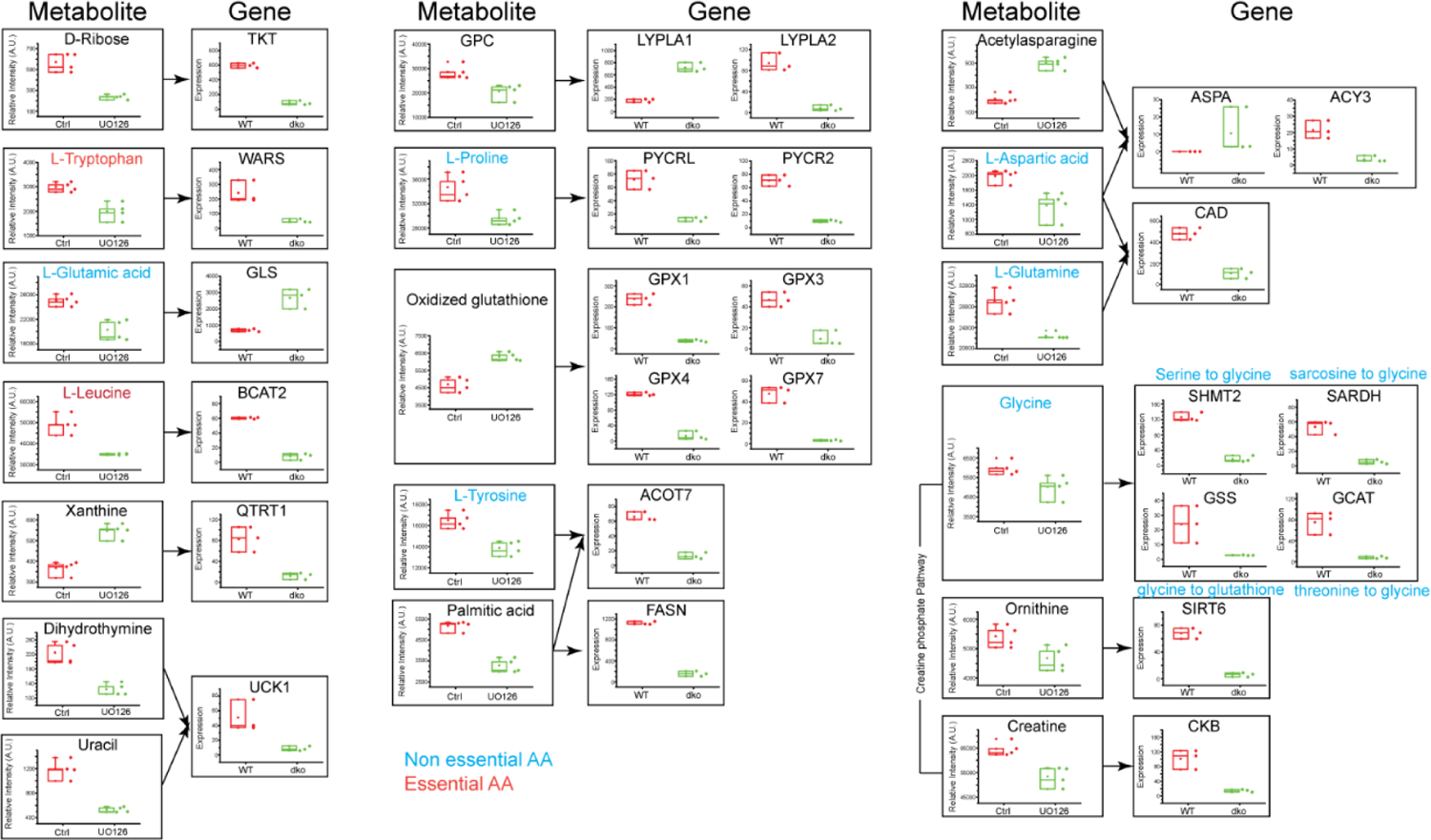
Cross linkage of identified metabolites and transcriptional changes caused by MAPK/ERK-deficiency. Quantitative metabolites and gene expression levels from metabolomics and RNAseq analysis were used to connect metabolite-gene expression correlations. Metabolite levels were measured by LC/MS from 4 independent control (DMSO treated) and four MEK-inhibited (UO126 treated) mK4 cell samples. Gene expression level changes were retrieved from RNAseq performed in isolated populations of three control and three *Mek1/2* double knockout NPs (Kurtzeborn et al, in revision).

The integrated systems biology approach identified several metabolic pathways affected by MAPK/ERK-deficiency. Among the 46 originally identified metabolites, 18 metabolites were correlated to 25 differentially expressed genes (Figure 4, Table 3) in our RNA-Seq data (Kurtzeborn et al, manuscript in revision). The most significant findings underline and confirm the severe shortage of energy, which is detected as diminished levels of ATP, ADP, Phosphoenolpyruvate, Ribose-5-phosphate, and 3-Phosphoglycerate and transcripts encoding their metabolic enzymes (Table 3). Interestingly, many of the diminished metabolic pathways were also related to amino acid metabolism. A few of these are tryptophan, shown to regulate pluripotent stem cell proliferation and expression of pluripotency regulator *Oct4*; glutamine, which regulates reversibility of hair follicle stem cells; and glycine, which is the major regulatory metabolite with functions in antioxidant response and neuronal specification (45–48). One additionally interesting metabolite identified here is proline, which not only regulates stem cell pluripotency but also contributes to the control of neuronal differentiation (38-40, 49-51). Proline metabolism is down-regulated in MAPK/ERK-deficient cells, and its direct regulators *Pycr2* (mitochondrial) and *Pycrl* (cytosolic) are also down-regulated in MAPK/ERK-deficient NPs. The correlations of metabolite-to-gene expression changes identified here are visualized in Figure 5.

**Figure 5.**
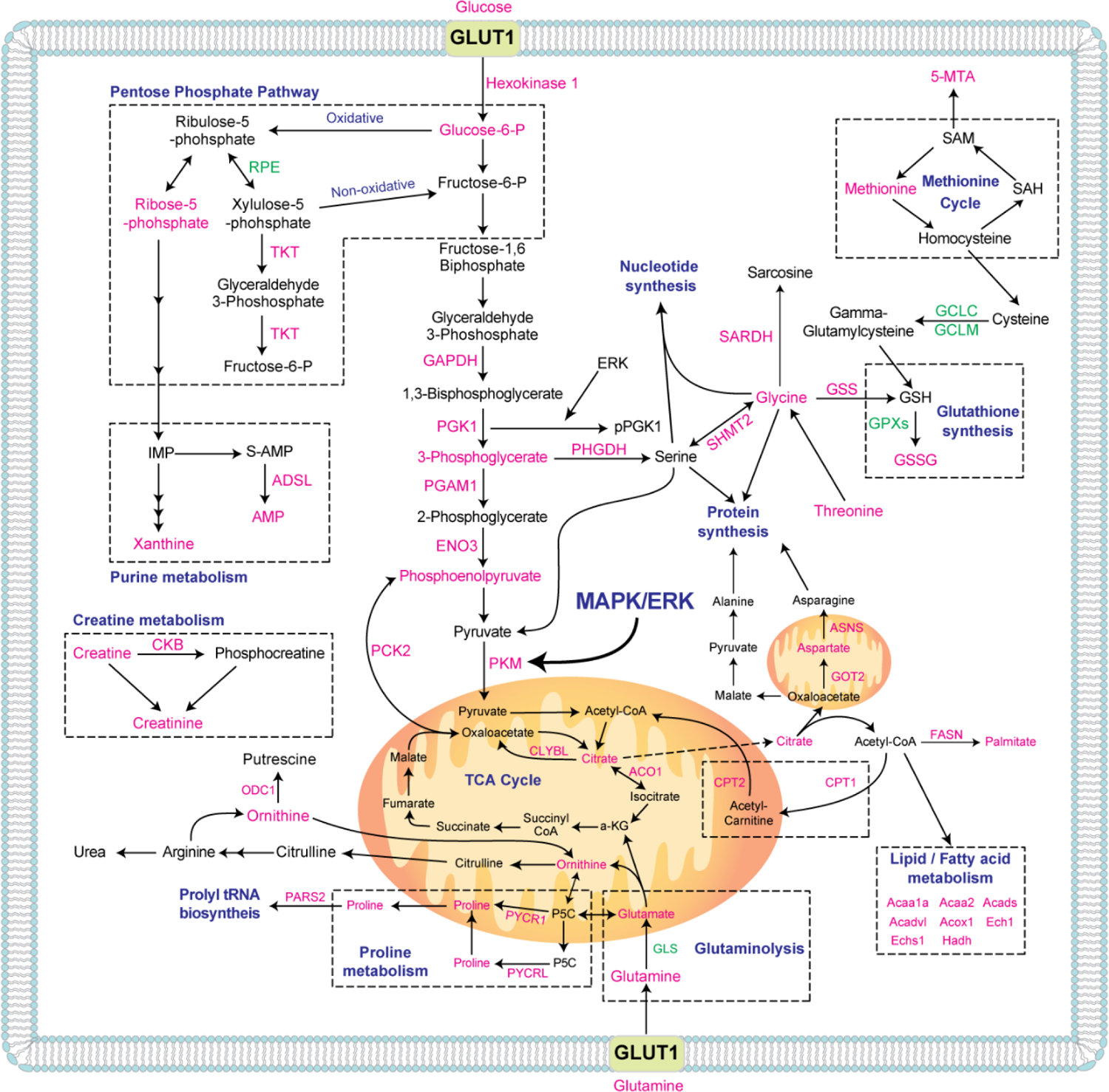
Presentation of overall pathway map of MAPK/ERK metabolic effects identified by integrating metabolomics and RNAseq data. Actual quantitated metabolites and enzymes are represented with magenta (down-regulated) and green (up-regulated). Dotted boxes represent metabolic pathways including metabolites and genes. RPE; Ribulose-phosphate 3-epimerase, TKT; Transketolase, GAPDH; Glyceraldehyde-3-phosphate dehydrogenase, PGK1; Phosphoglycerate kinase 1, PHGDH; D-3-phosphoglycerate dehydrogenase, PGAM1; Phosphoglycerate mutase 1, ENO3; Beta-enolase, PCK2; Phosphoenolpyruvate carboxykinase [GTP], PKM; Pyruvate kinase, ADSL; Adenylosuccinate lyase, CKB; Creatine kinase B-type, ODC1; Ornithine decarboxylase, PARS2; Probable proline--tRNA ligase, PYCR1; pyrroline-5-carboxylate reductase 1, PYCRL; Pyrroline-5-carboxylate reductase, GLS; Glutaminase, CLYBL; Citramalyl-CoA lyase, ACO1; Aconitate hydratase 1, CPT1; Carnitine palmitoyltransferase 1, CPT2; Carnitine palmitoyltransferase 2, FASN; Fatty Acid Synthase, GCLC; Glutamate--cysteine ligase catalytic subunit, GCLM; Glutamate--cysteine ligase regulatory subunit, SARDH; Sarcosine dehydrogenase, SHMT2; Serine hydroxymethyltransferase 2, GSS; Glutathione synthetase, GPx; Glutathione peroxidase, ASNS; Asparagine synthetase, GOT2; Aspartate aminotransferase. Magenta represents down-regulated, green represents up-regulated.

**Table 3.**
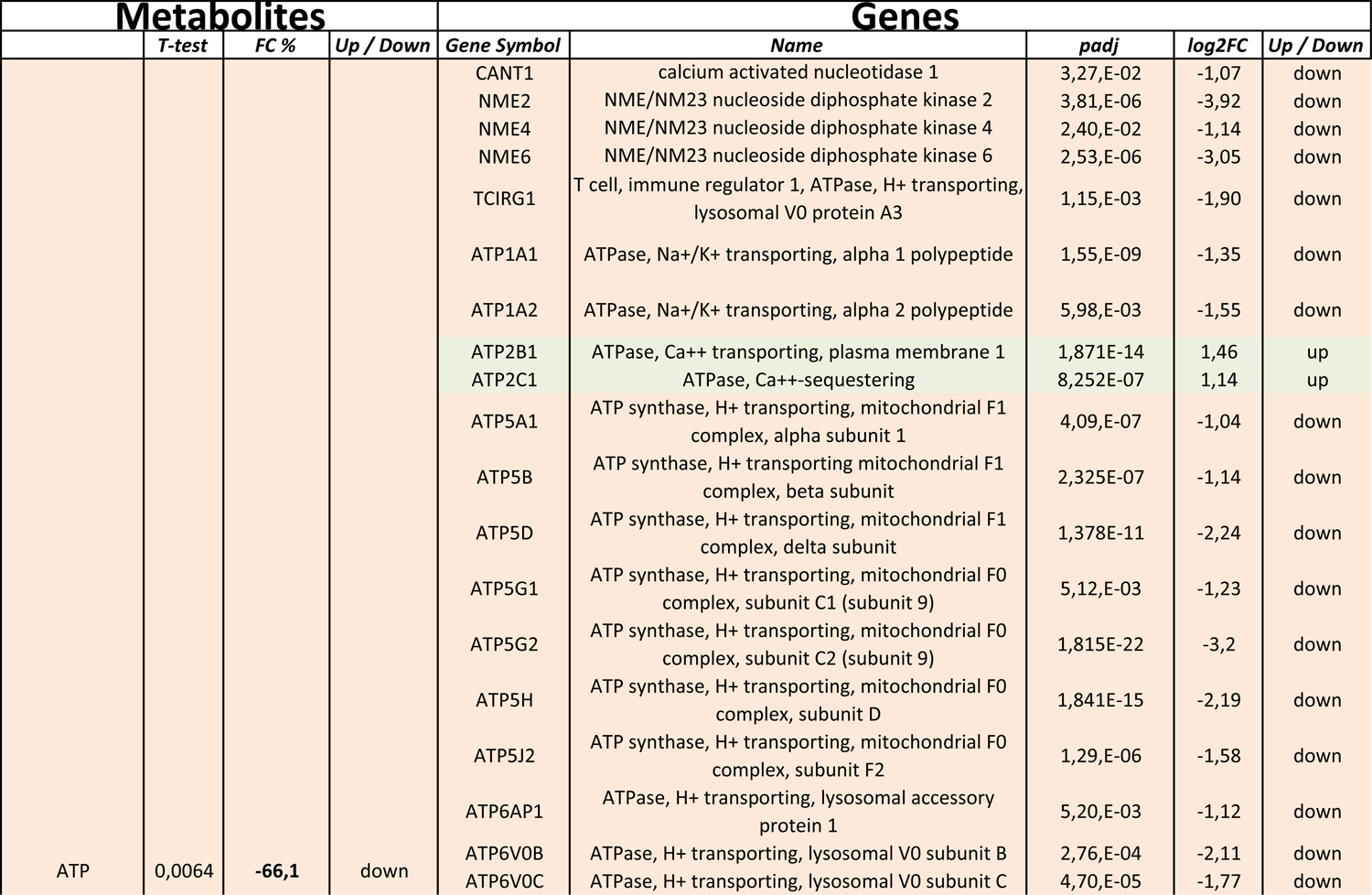
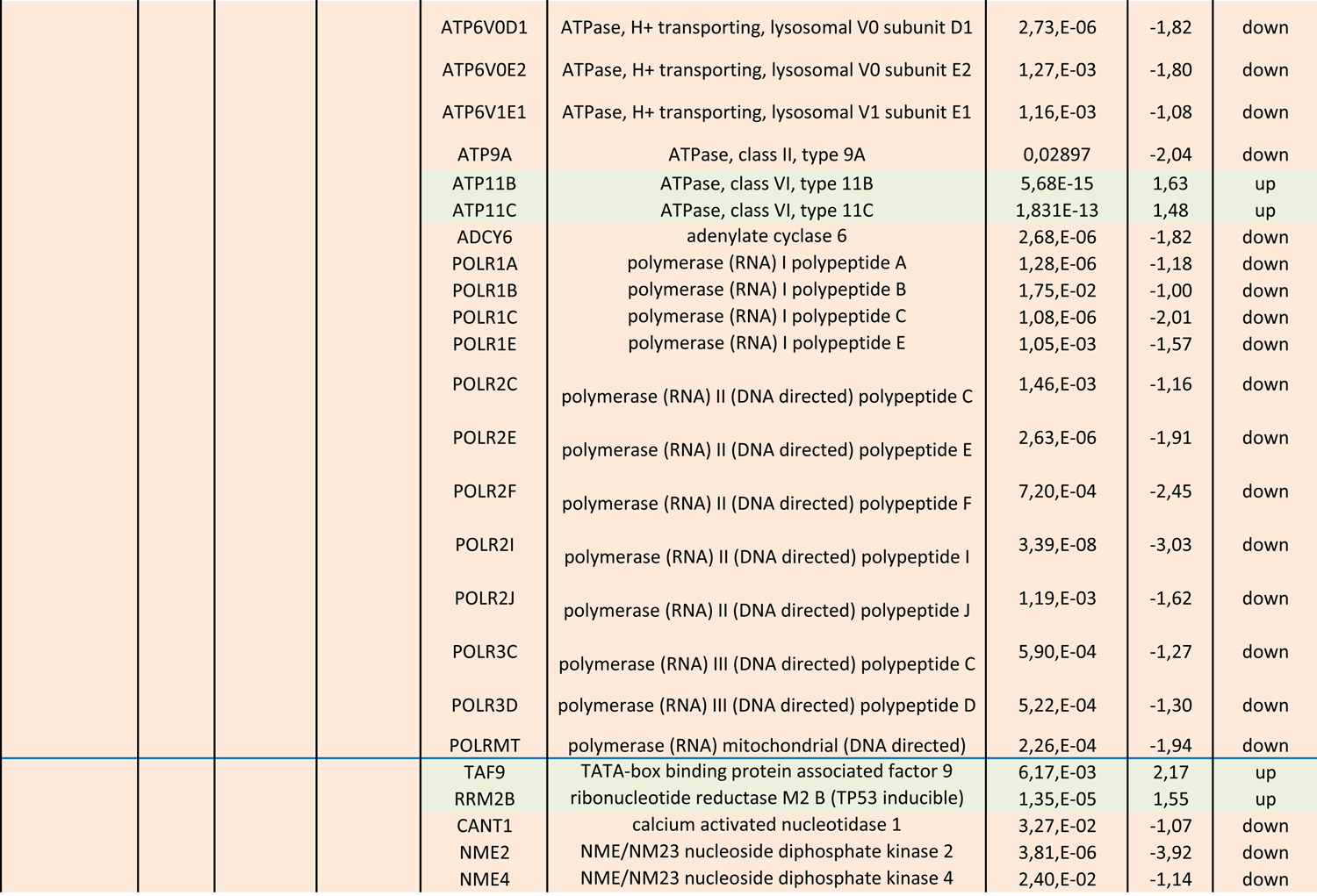
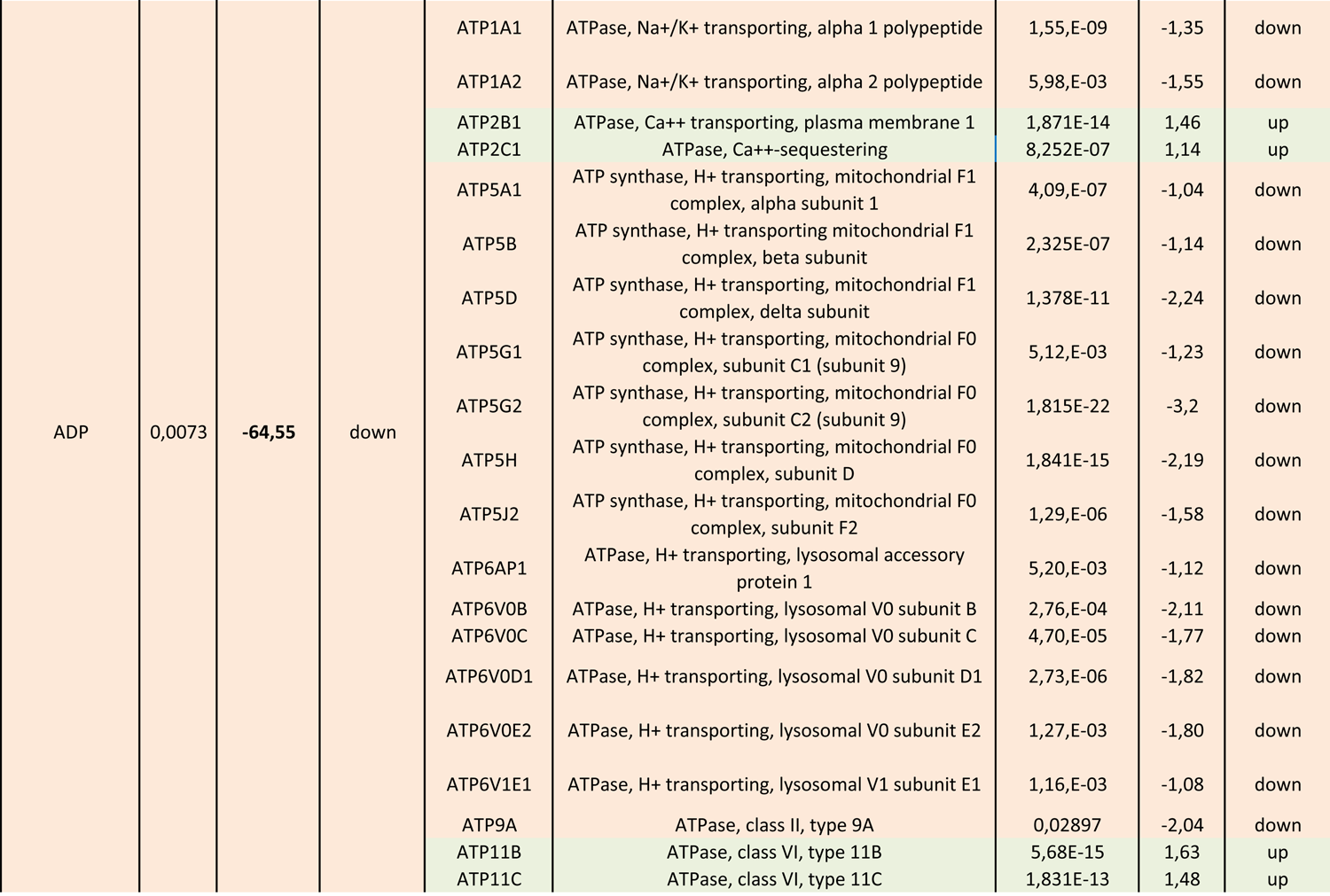
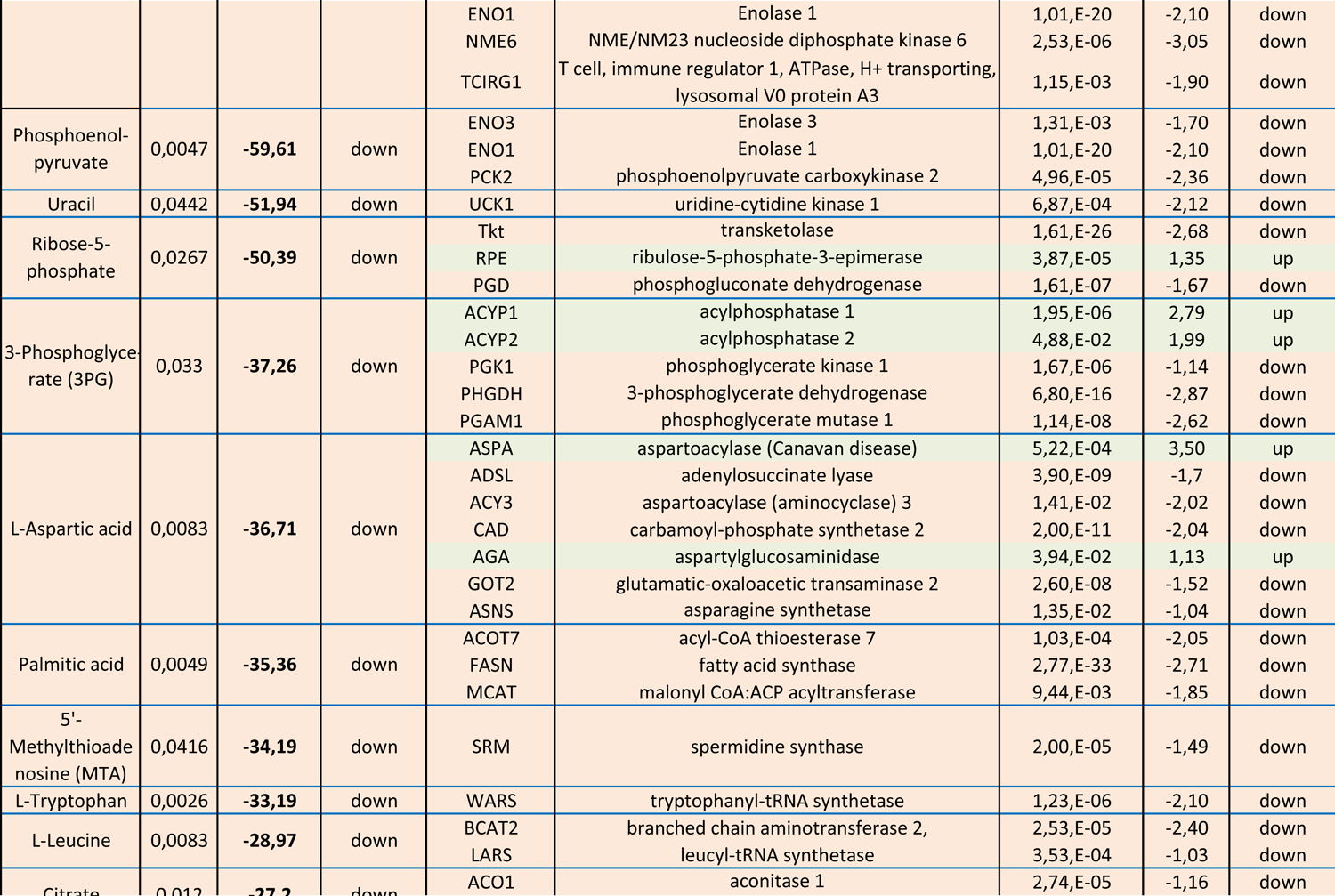
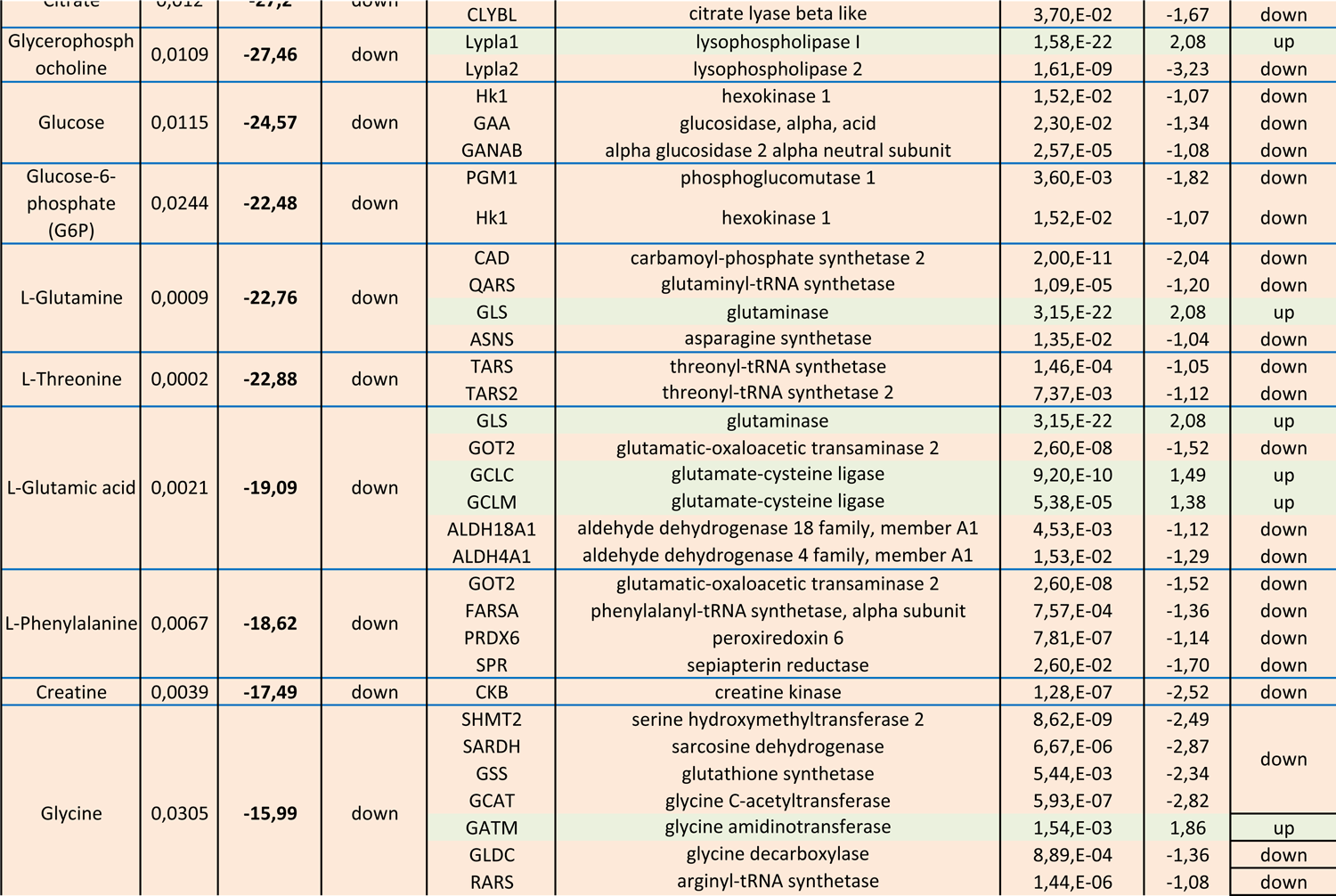
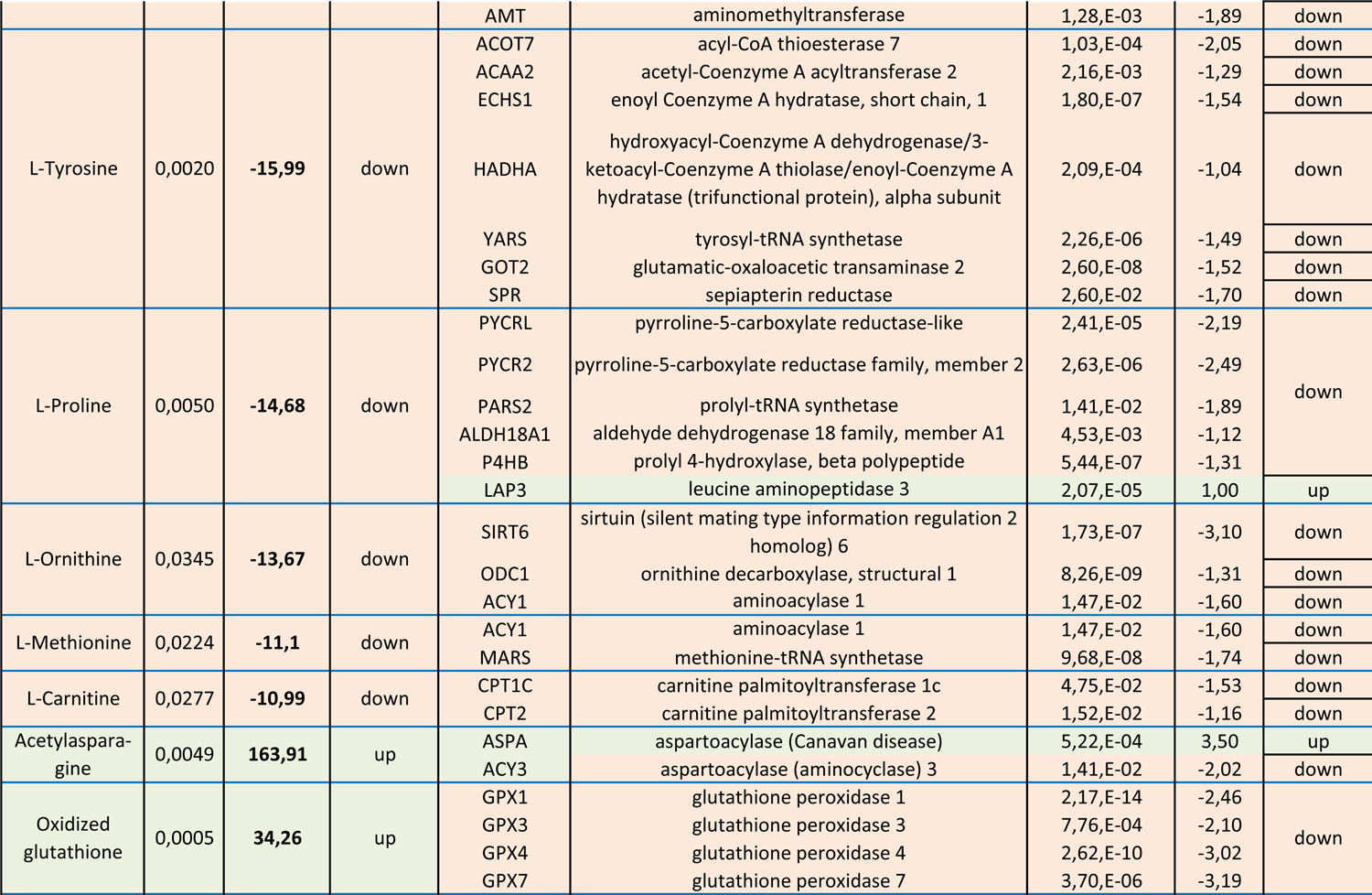
The correlations between identified metabolites and differentially expressed genes

### Pyruvate dose-dependently affects nephron progenitor niche morphology

To investigate the physiological consequences of identified metabolic changes, we first assayed the effect of pyruvate on early kidney development. Embryonic day 11.5 (E11.5) kidneys were cultured for 72h *in vitro* as described earlier (52) under normal (1mM) and different pyruvate concentration conditions. Cultured kidneys were immunostained with calbindin to visualize ureteric bud tips, which form an extrinsic niche for NPs (10), and branching morphogenesis (Figure 6A). Without pyruvate, kidney growth through branching morphogenesis was severely compromised (50-80% less tips), the ratio of ureteric bud tips-to-trunks was obscured, and the tips were enlarged, especially with 0.5mM pyruvate concentrations. Pyruvate supplementation with low doses, concentrations less than 1.0 mM, improved the branching rate and pattern but branching was not completely rescued to the level typical for control kidneys (1.0 mM pyruvate). Interestingly, pyruvate concentration higher than 1mM resulted in diminished branching and tip amounts but with a different pattern than in low concentrations (0-0.5mM) and without a dramatic effect on tip morphology.

**Figure 6.**
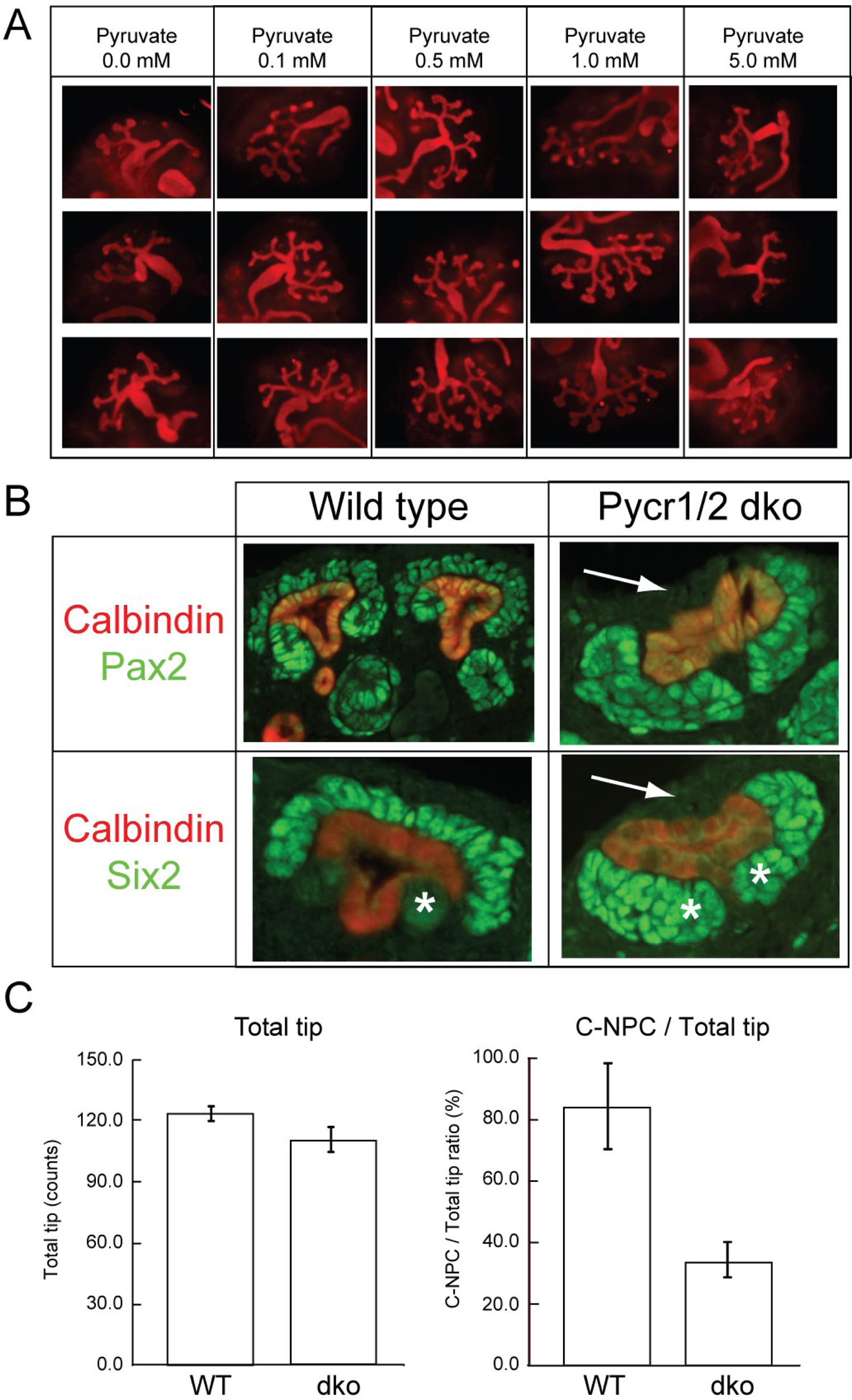
Functional characterization of identified metabolite effects on nephron progenitor biology. **(A)** Embryonic kidneys at E11.5 were cultured for 72 hours under different concentration of pyruvate supplementation, and immunostained against Calbindin, which visualizes the entire ureteric bud epithelium. Normal kidney culture medium contains 1mM pyruvate, which serves as a control for other concentrations. (**B**) Postnatal day 1 control (wild type) and *Pycr1/2* double knockout (dko) kidneys were immunostained for PAX2 (green, marker of nephron progenitors and precursors) or SIX2 (green, marker of nephron progenitors) in combination with Calbindin (red, marker of ureteric bud epithelium). White arrows indicate the regions where PAX2- and SIX2-positive NP cells should be located but are significantly diminished in the absence of *Pycr1/2*. White asterisks point to the differentiating nephron precursors, which show only very weak SIX2 signal in control kidneys but to where SIX2 almost exclusively localizes in *Pycr1/2* dko kidneys. (C) Quantification of ureteric tip numbers (total tip) and cortical nephron progenitors (C-NPC) per each tip in the cortex of wild type and *Pycr1/2* dko kidneys (n=2 kidney/ genotype, total of 15 randomly selected medullar sections were counted).

### Proline metabolism is required for nephron progenitor maintenance

Our integrated metabolomics and transcriptomics data revealed that the majority of pathways involved in energy metabolism are down-regulated upon MAPK/ERK-deficiency. Down-regulation of proline metabolism is of particular interest as in addition to reduction of the metabolite itself, the expression of its regulatory genes *Pycr2* and *Pycrl* was significantly diminished (Figure 4).

To assess the physiological *in vivo* requirement of proline metabolism in kidney development, we utilized conventional inactivation of *Pycr1* and -*2* genes in mouse. We found that inactivation of *Pycr* genes resulted in Mendelian ratios of homozygous *Pycr1^-/-^*, *Pycr2^-/-^* and double knockout *Pycr1^-/-^;Pycr2^-/-^* pups with grossly normal kidney morphology and histology (Supplementary Figure 1). Further characterization of *Pycr1^-/-^;Pycr2^-/-^* mouse kidneys at postnatal day 1 (P1) revealed defects in NP population as detected by a reduced number of PAX2-positive cells per ureteric bud and by premature differentiation of postnatal SIX2-positive NPs (Figure 6B). These results demonstrate that proline metabolism, which we identified here to depend on MAPK/ERK activation, is essential for the maintenance of NP population in the developing kidney

## DISCUSSION

Metabolomics in combination with systems biology is powerful approach to study functional linkages of metabolic and transcriptional changes (53, 54). Metabolomics has the potential to link genotype to phenotype (55), and it has been broadly and successfully used in systems biology studies (56) to gain a better understanding of complex biological systems, as exemplified by metabolomics studies of mitochondrial dysfunction in kidney diseases (57–61). We have previously demonstrated the importance of MAPK/ERK pathway in NP maintenance and differentiation during embryonic kidney development (29). Here we combined metabolomics and transcriptomics characterization of MAPK/ERK-deficient NP cells to advance the understanding of cellular defects contributing to congenital kidney anomalies.

Congenital anomalies of the kidney and urinary tract (CAKUT) are common birth defects caused by abnormalities in kidney morphogenesis, which contribute to the growing worldwide chronic kidney disease burden (62–64). The spectrum of CAKUT phenotypes varies from mild renal hypoplasia with reduced nephron count, to hypodysplasia and a complete lack of kidneys, a condition known as agenesis or aplasia (65). The milder CAKUT forms such as reduced nephron mass due to impaired NP maintenance or differentiation are not life-threatening at early ages but do possess increased risk of chronic kidney disease with cardiovascular co-morbidities later in life (1, 2). As the final nephron count in each individual is determined by renal morphogenesis during fetal development, any factor possibly affecting the rate and extent of nephrogenesis is of high interest.

There is increasing awareness of how energy metabolism and especially mitochondrial dysfunction contributes to various congenital, acute, and chronic pathologies. Cancer cells are known for their aerobic glycolysis, a.k.a Warburg effect, during which glucose uptake and its conversion to lactate are high and a similar metabolic profile is recognized in actively proliferating cells including stem cells and fetal neuronal tissues (66, 67). MAPK/ERK pathway regulates ATP levels and energy availability through the control of α-enolase, which is a glycolytic pyruvate producing enzyme (68). Adult kidney is a highly energetic organ that contains abundant mitochondria (69) and where malfunctioning mitochondrial energy metabolism has significant effects on the pathogenesis of adult-onset kidney diseases (70). Therefore, mitochondria have emerged as promising therapeutic options for renal diseases (71, 72).

Most metabolite-gene interactions observed here showed the same change patterns presenting decreased measures for both. For example, we identified that the proline production-related enzymes PYCRL and PYCR2 were down-regulated, and their metabolic product, L-proline, was also diminished. As another example, we identified a significant decrease in both acid synthase (FASN) which is required to produce its end-metabolite product palmitic acid. Conversely, in a few cases, the correlation between the metabolite level and the gene expression was opposite to each other. For instance, subtypes of glutathione peroxidase family were down-regulated (GPX1, GPX3, GPX4, and GPX7), but the oxidized glutathione produced by these genes was increased demonstrating that transcriptional changes do not always translate into protein level differences. All the possible correlations between identified metabolites and DEGs are summarized in Table 3.

Our integrated metabolomics and RNA-Seq characterization of MAPK/ERK-deficient NPs revealed significant gene expression changes in mitochondrial energy metabolism (30%) including oxidative phosphorylation (19%) and mitochondrion (11%). The majority of these transcripts and metabolites were downregulated by MAPK/ERK deficiency. Our experiments revealed diminished mitochondrial counts, reduced expression of α-enolase and decreased ATP levels in NPs with MAPK/ERK-deficiency. These likely stem from the block in glycolysis, as seen by diminished phosphoenolpyruvate (PEP) levels in the absence of MAPK/ERK activation. Under normal conditions, mitochondrial oxaloacetate can be metabolized to PEP by phosphoenolpyruvate carboxykinase 2 (PCK2), but as we also observed decreased expression of *Pck2* upon MEPK/ERK-deficiency this supplementation pathway likely is unable to rescue reduced PEP levels. These results collectively demonstrated that the MAPK/ERK activity in NPs significantly contributes to cellular metabolism, particularly to mitochondrial energy metabolism.

In addition to energy metabolism, we identified a shortage of precursors for nucleotide and protein biosynthesis. Maintenance of the NP population throughout embryonic and early postnatal periods depends on their persistent proliferation that replenishes induced NPs destined for differentiation (10). We have previously reported that MAPK/ERK deficiency causes NP proliferation decline due to diminished DNA replication, which contributes to the failure in their maintenance (29). Glucose metabolism by glycolysis and mitochondrial tricarboxylic acid (TCA) cycle are recognized as fundamental bases for DNA synthesis and post-translational modifications of proteins and nucleotides (16, 73). Their disturbance in the absence of MAPK/ERK activation contributes to the failure to maintain NP population *in vivo*.

Production of pyruvate, one of the major metabolite precursors playing essential functions in mitochondrial metabolism, requires function of the MAPK/ERK-regulated glycolytic enzyme pyruvate kinase isozyme M2 (PKM2), which is expressed by the embryonic kidney (23, 74). In adult kidneys, PKM2 is shown to be protective against acute injury (24). Our results demonstrated decreased expression of *Pkm2* in MAPK/ERK-deficient NPs and reduced levels of TCA intermediate citrate. Analysis of the effects of different pyruvate concentrations in kidney cultures showed dose-dependent changes in the morphology of ureteric bud tips, which serve as an extrinsic niche for nephron progenitors (10). The experiment suggests that low pyruvate availability, for example due to diminished conversion of precursor PEP to pyruvate, supports ureteric bud tip expansion but reduces the number of tip niches, which are shown to have detrimental effects on NP maintenance (12). High pyruvate concentrations also reduce tip niche numbers but do not significantly affect tip morphology. Interestingly, forced pyruvate redirecting to mitochondrial acetyl-CoA production increases pluripotency while glycolytic Ac-CoA production drives histone acetylation in embryonic stem cells (22) suggesting that pyruvate availability critically contributes to the NP stemness.

We also identified down-regulation of *Pycr2* and *Pycrl* genes, which regulate the proline in cells and demonstrated reduced levels of proline metabolite in MAPK/ERK-deficient cells. Proline metabolism has versatile functions in cellular homeostasis (36) and its role as a stem cell regulator is of high relevance to our study. Proline supplementation to embryonic stem cells induces their proliferation and induction towards epiblast while maternal proline improves nutrient transport to better support pre-implantation embryo development (38, 49–51). Proline deficiency causes several different inherited diseases that typically affect growth, central nervous system and renal functions (36). Our analysis of proline deficiency caused by simultaneous inactivation of *Pycr1* and *Pycr2* in developing kidney revealed its importance for NP maintenance in postnatal kidney. This is an important result not only because the exact functions of proline metabolism at organ and tissue level remained unknown, but also due to its therapeutic potential as a metabolite that is easy to administer.

## MATERIALS AND METHODS

### RNA isolation for sequencing, and data analysis

For sequencing, RNA was isolated from FAC-sorted nephron progenitors at E13.5 by a standard protocol for chloroform/isopropanol extraction, followed by DNase I treatment according to manufacturer’s instructions (Thermo Fisher Scientific – 04/2016, rev. B.00). Total RNA quality and quantity were estimated by using the Bioanalyzer RNA Total Pico (Agilent Technologies, Inc.) and Nanodrop analysis. Library preparation was done using NuGen Ovation Solo and sequencing was performed with NextSeq at BIDGEN DNA Sequencing and Genomics Laboratory (University of Helsinki). bcl2fastq2 Conversion Software was used to convert BCL files to FASTQ file format and demultiplex samples. Sequenced reads were trimmed for adaptor sequence and masked for low-complexity or low-quality sequence using Trimmomatic. Trimmed reads were mapped to GENCODE Mus musculus Release M23 reference genome GRCm38 using STAR aligner (2.6.0c).

Raw gene counts were normalized using DESeq2 R-package (v.3.6.2) and sample clustering was visualized by PCA-plots. Multiple testing adjustment of p-values was done with Benjamini-Hochberg procedure to compare gene expressions in three control and three nephron-specific MAPK/ERK-deficient samples (*Six2-TGC;Mek1^fl/fl^;Mek2^-/-^*).

### Gene Ontology and Pathway Analysis

To further analyze how MAPK/ERK-deficient nephron progenitor (NP) populations have different metabolic characteristics, we utilized our recent whole-genome RNA-Seq data (Kurtzeborn et al, manuscript in revision). To interpret differentially expressed genes in NPs, we performed the pathway analysis using Cytoscape plug-in ClueGO to create a functionally organized gene ontology (GO). The GO_Cellular Component and KEGG ontology were used for GO analysis and we performed the GO analysis using total DEGs and up-regulated and down-regulated DEGs separately.

### Metabolomics sample preparation

To investigate metabolic changes in NPs incurred by MEK inhibition, we analyzed metabolic changed of mK4 cell line, derived from embryonic kidney mesenchyme, through a LC/MS-based metabolomics approach. Prior to cell harvest for metabolite extraction, mK4 cells were incubated with DMSO (0.1% v/v), MEK inhibitor U0126 (15 µM), or without any treatment. After 24 hours incubation, cells were harvested using 0.25% Trypsin/EDTA treatment and washed with cold PBS. The metabolites were extracted by the same method applied in our current RNA-Seq research (Kurtzeborn et al, manuscript in revision). In brief, harvested cells were dissolved with 600 µl of methanol/chloroform solvent (2:1 v/v ratio), quick frozen twice with liquid nitrogen and thawed at room temperature. Then, 200 µl of each of chloroform and distilled water were sequentially added, and metabolites were separated by following centrifugation at 15,000 g for 20 min at 4 °C. Finally, water soluble metabolites (upper phase after centrifugation) were dried using centrifugal vacuum evaporator. The middle phase (protein aggregate) after centrifugation was isolated and dissolved in 6M urea buffer and total protein concentration were measured to utilize for normalization.

### Liquid chromatography-mass spectrometry (LC-MS) analysis

The ZIC-HILIC column (3.5 µm, 200 Å, 100 × 2.1 mm; Merck Millipore) and Acquity UPLC system (Waters, MA, USA) were used for liquid chromatography. The Q Exactive Focus Orbitrap Mass Spectrometer (Thermo Fisher Scientific) equipped with HESI sources was used for metabolite identification. Chromatographic separation was performed on a ZIC-HILIC column (3.5 µm, 200 Å, 100 × 2.1 mm; Merck Millipore) using an Acquity UPLC system (Waters, MA, USA). The column temperature was 35°C with a flow rate of 0.4 mL/min and the auto-sampler cooler temperature was set at 4°C with an injection volume of 2 µl. Analytes were eluted with a mobile phase composed of 10 mM ammonium acetate in water with pH 6.8 (A) and 10 mM ammonium acetate in water:acetonitrile solution (1:3) with pH 6.8 (B). Gradient conditions were as follows: 0–1 min kept B as 100%, 1–7 min declined B from 100% to 20%, 7–11 min kept B as 20%, 11.1 min set B as 100% and kept the composition until 23 min. Mass spectrometry experiments were performed on a Q Exactive Focus Orbitrap Mass Spectrometer (Thermo Fisher Scientific) equipped with HESI sources. To detect as many ions as possible, both positive and negative detection modes were used in the LC/MS experiment. The inclusion list for parallel reaction method is shown in supplementary Table S2. The MS2 spectrum patterns comparison to the online MS database Metlin and high-resolution m/s from full MS experiment were applied to confirm target metabolites. All the measured data were normalized against total protein concentration which were measured using middle phase protein aggregate concentration.

### Metabolomics analysis and systems biology approach

The multivariate statistical analysis was carried out using SIMCA-P version 11.0 program (Umetrics, Umeå, Sweden). To distinguish metabolic pattern differences from each experimental group, the supervised statistical method, partial least squares discriminant analysis (PLS-DA) analysis, was performed using the processed data from LC/MS positive and negative modes, and statistically contributing metabolites on group separation on PLS-DA were identified. To further analyze the identified metabolites, we performed Metabolite Set Enrichment Analysis (MSEA) thorough web-based analysis tool provided from MetaboAnalyst (75), and the levels of individual contributing metabolites on MSEA result were visualized by heatmap analysis in R package.

### Pyruvate effect on embryonic kidney culture

E11.5 control mouse kidneys were cultured under different concentrations of pyruvate supplementation for 72 hours. Cultured embryonic kidneys were analyzed via immunostaining with CALBINDIN, which labels ureteric bud epithelium and allows to determine the effect of pyruvate concentration on ureteric bud branching.

### Hematoxylin-eosin staining, immunofluorescent detection of nephron progenitors and precursors, imaging and quantification

Postnatal day 1 (P1) kidneys from Pycr1/2 mating were collected and fixed with 4% PFA, after which they were processed to paraffin and sectioned according to standard procedures. Hematoxylin-eosin and immunofluorescent staining, and imaging procedures were performed as previously described (20). Primary antibodies used are CALBINDIN (Santa Cruz Biotechnology, sc-7691) PAX2 (Invitrogen, 71-6000) and SIX2 (ProteinTech, 11562-1-AP). Alexa Fluor-conjugated secondary antibodies (1:400; Jackson Immuno Research Laboratories) were used as secondary antibodies. Quantification of nephron progenitor cells, ureteric tips and T-buds was performed blindly by counting 15 medullar sections in two wild type and two double knockout *Pycr1/2* kidneys by three independent researchers. Mean of three counts were used to calculate mean value of cortical nephron progenitors per each ureteric bud tip.

## Author Contributions

H.N.K: Research design, experiment, analysis data, manuscript writing; K.K: Experiment, manuscript editing; X.J.: LC/MS experiment; S.P.: Manuscript editing; B.R., A.L and N.E.B mouse *Pycr1/2* knockouts, S.K.: Conceptualization of the research, manuscript writing and editing.

## References

1. Bertram JF, Douglas-Denton RN, Diouf B, Hughson MD, & Hoy WE (2011) Human nephron number: implications for health and disease. Pediatr. Nephrol. 26(9):1529–1533.

2. Luyckx VA & Brenner BM (2010) The clinical importance of nephron mass. J. Am. Soc. Nephrol. 21(6):898–910.

3. Davidson AJ, Lewis P, Przepiorski A, & Sander V (2019) Turning mesoderm into kidney. *Semin. Cell Dev. Biol.*, (Elsevier), pp 86–93.

4. Kurtzeborn K, Kwon HN, & Kuure S (2019) MAPK/ERK Signaling in Regulation of Renal Differentiation. Int. J. Mol. Sci. 20(7):1779.

5. Costantini F & Kopan R (2010) Patterning a complex organ: branching morphogenesis and nephron segmentation in kidney development. Dev. Cell 18(5):698–712.

6. Oxburgh L (2018) Kidney Nephron Determination. Annu Rev Cell Dev Biol 34:427–450.

7. Hartman HA, Lai HL, & Patterson LT (2007) Cessation of renal morphogenesis in mice. Dev Biol 310(2):379–387.

8. Hinchliffe SA, Sargent PH, Howard CV, Chan YF, & van Velzen D (1991) Human intrauterine renal growth expressed in absolute number of glomeruli assessed by the disector method and Cavalieri principle. Lab Invest 64(6):777–784.

9. Ryan D, et al. (2018) Development of the Human Fetal Kidney from Mid to Late Gestation in Male and Female Infants. EBioMedicine 27:275–283.

10. O’Brien LL (2019) Nephron progenitor cell commitment: Striking the right balance. *Semin. Cell Dev. Biol.*, (Elsevier), pp 94–103.

11. Li H, Hohenstein P, & Kuure S (2021) Embryonic Kidney Development, Stem Cells and the Origin of Wilms Tumor. Genes (Basel) 12(2).

12. Li H, et al. (2021) Postnatal prolongation of mammalian nephrogenesis by excess fetal GDNF. Development 148(10).

13. Brown AC, Muthukrishnan SD, & Oxburgh L (2015) A synthetic niche for nephron progenitor cells. Dev. Cell 34(2):229–241.

14. Volovelsky O, et al. (2018) Hamartin regulates cessation of mouse nephrogenesis independently of Mtor. Proc Natl Acad Sci U S A 115(23):5998–6003.

15. Cargill K & Sims-Lucas S (2020) Metabolic requirements of the nephron. Pediatr. Nephrol. 35(1):1–8.

16. Tortelote GG, Colon-Leyva M, & Saifudeen Z (2021) Metabolic programming of nephron progenitor cell fate. Pediatr. Nephrol. 36(8):2155–2164.

17. Liu J, et al. (2017) Regulation of Nephron Progenitor Cell Self-Renewal by Intermediary Metabolism. J. Am. Soc. Nephrol. 28(11):3323–3335.

18. Shyh-Chang N & Ng HH (2017) The metabolic programming of stem cells. Genes Dev. 31(4):336–346.

19. Cargill K, et al. (2019) Von Hippel-Lindau Acts as a Metabolic Switch Controlling Nephron Progenitor Differentiation. J Am Soc Nephrol 30(7):1192–1205.

20. Conaghan J, Handyside AH, Winston RM, & Leese HJ (1993) Effects of pyruvate and glucose on the development of human preimplantation embryos in vitro. J. Reprod. Fertil. 99(1):87–95.

21. Butcher L, Coates A, Martin KL, Rutherford AJ, & Leese HJ (1998) Metabolism of pyruvate by the early human embryo. Biol. Reprod. 58(4):1054–1056.

22. Moussaieff A, et al. (2015) Glycolysis-mediated changes in acetyl-CoA and histone acetylation control the early differentiation of embryonic stem cells. Cell Metab. 21(3):392–402.

23. Yang W, et al. (2012) ERK1/2-dependent phosphorylation and nuclear translocation of PKM2 promotes the Warburg effect. Nat. Cell Biol. 14(12):1295–1304.

24. Zhou H-L, et al. (2019) Metabolic reprogramming by the S-nitroso-CoA reductase system protects against kidney injury. Nature 565(7737):96–100.

25. Magadum A, et al. (2020) Pkm2 Regulates Cardiomyocyte Cell Cycle and Promotes Cardiac Regeneration. Circulation 141(15):1249–1265.

26. Purcell NH, et al. (2007) Genetic inhibition of cardiac ERK1/2 promotes stress-induced apoptosis and heart failure but has no effect on hypertrophy in vivo. Proceedings of the National Academy of Sciences 104(35):14074–14079.

27. Maejima Y, Galeotti J, Molkentin JD, Sadoshima J, & Zhai P (2012) Constitutively active MEK1 rescues cardiac dysfunction caused by overexpressed GSK-3α during aging and hemodynamic pressure overload. American Journal of Physiology-Heart and Circulatory Physiology 303(8):H979–H988.

28. Ihermann-Hella A, et al. (2014) Mitogen-activated protein kinase (MAPK) pathway regulates branching by remodeling epithelial cell adhesion. PLoS Genet. 10(3):e1004193.

29. Ihermann-Hella A, et al. (2018) Dynamic MAPK/ERK Activity Sustains Nephron Progenitors through Niche Regulation and Primes Precursors for Differentiation. Stem Cell Reports 11(4):912–928.

30. Li Y, et al. (2015) p53 Enables metabolic fitness and self-renewal of nephron progenitor cells. Development 142(7):1228–1241.

31. Kimura T, et al. (2003) Impaired function of p53R2 in Rrm2b-null mice causes severe renal failure through attenuation of dNTP pools. Nat Genet 34(4):440–445.

32. Powell DR, et al. (2005) Rapid development of glomerular injury and renal failure in mice lacking p53R2. Pediatr Nephrol 20(3):432–440.

33. Kuo ML, et al. (2016) PYCR1 and PYCR2 Interact and Collaborate with RRM2B to Protect Cells from Overt Oxidative Stress. Sci Rep 6:18846.

34. Aguilera O, et al. (2016) Vitamin C uncouples the Warburg metabolic switch in KRAS mutant colon cancer. Oncotarget 7(30):47954–47965.

35. de Koning TJ (2017) Amino acid synthesis deficiencies. J Inherit Metab Dis 40(4):609–620.

36. Patriarca EJ, et al. (2021) The Multifaceted Roles of Proline in Cell Behavior. Front Cell Dev Biol 9:728576.

37. D’Aniello C, et al. (2017) Vitamin C and l-Proline Antagonistic Effects Capture Alternative States in the Pluripotency Continuum. Stem Cell Reports 8(1):1–10.

38. Cermola F, et al. (2021) Gastruloid Development Competence Discriminates Different States of Pluripotency. Stem Cell Reports 16(2):354–369.

39. Escande-Beillard N, et al. (2020) Loss of PYCR2 Causes Neurodegeneration by Increasing Cerebral Glycine Levels via SHMT2. Neuron 107(1):82–94 e86.

40. Stum MG, et al. (2021) Genetic analysis of Pycr1 and Pycr2 in mice. Genetics 218(1).

41. Bindea G, et al. (2009) ClueGO: a Cytoscape plug-in to decipher functionally grouped gene ontology and pathway annotation networks. Bioinformatics 25(8):1091–1093.

42. Shannon P, et al. (2003) Cytoscape: a software environment for integrated models of biomolecular interaction networks. Genome Res. 13(11):2498–2504.

43. Ashburner M, et al. (2000) Gene ontology: tool for the unification of biology. Nat. Genet. 25(1):25–29.

44. Kanehisa M & Goto S (2000) KEGG: kyoto encyclopedia of genes and genomes. Nucleic Acids Res. 28(1):27–30.

45. Avila A, et al. (2014) Glycine receptors control the generation of projection neurons in the developing cerebral cortex. Cell Death Differ. 21(11):1696–1708.

46. Cheng J, et al. (2015) Tryptophan derivatives regulate the transcription of Oct4 in stem-like cancer cells. Nat Commun 6(1):7209.

47. Kim CS, et al. (2020) Glutamine metabolism controls stem cell fate reversibility and long-term maintenance in the hair follicle. Cell Metab. 32(4):629–642. e628.

48. Someya S, et al. (2021) Tryptophan Metabolism Regulates Proliferative Capacity of Human Pluripotent Stem Cells. iScience 24(2):102090.

49. Washington JM, et al. (2010) L-Proline induces differentiation of ES cells: a novel role for an amino acid in the regulation of pluripotent cells in culture. Am. J. Physiol. Cell Physiol. 298(5):C982–992.

50. Casalino L, et al. (2011) Control of embryonic stem cell metastability by L-proline catabolism. J. Mol. Cell. Biol. 3(2):108–122.

51. Liu N, et al. (2019) Maternal L-proline supplementation enhances fetal survival, placental development, and nutrient transport in mice. Biol. Reprod. 100(4):1073–1081.

52. Ihermann-Hella A & Kuure S (2019) Mouse ex vivo kidney culture methods. Kidney Organogenesis, (Springer), pp 23-30.

53. Pang H, Jia W, & Hu Z (2019) Emerging applications of metabolomics in clinical pharmacology. Clin. Pharmacol. Ther. 106(3):544–556.

54. Kitano H (2002) Systems biology: a brief overview. Science 295(5560):1662–1664.

55. Fiehn O (2002) Metabolomics--the link between genotypes and phenotypes. Plant Mol. Biol. 48(1-2):155–171.

56. Weckwerth W (2003) Metabolomics in systems biology. Annu. Rev. Plant Biol. 54:669–689.

57. Nam H, Chung BC, Kim Y, Lee K, & Lee D (2009) Combining tissue transcriptomics and urine metabolomics for breast cancer biomarker identification. Bioinformatics 25(23):3151–3157.

58. McKillop AM & Flatt PR (2011) Emerging applications of metabolomic and genomic profiling in diabetic clinical medicine. Diabetes Care 34(12):2624–2630.

59. Atzler D, Schwedhelm E, & Zeller T (2013) Integrated genomics and metabolomics in nephrology. Nephrology Dialysis Transplantation 29(8):1467–1474.

60. Sharma K, et al. (2013) Metabolomics reveals signature of mitochondrial dysfunction in diabetic kidney disease. J. Am. Soc. Nephrol. 24(11):1901–1912.

61. Tran MT, et al. (2016) PGC1α drives NAD biosynthesis linking oxidative metabolism to renal protection. Nature 531(7595):528–532.

62. Luyckx VA, Tonelli M, & Stanifer JW (2018) The global burden of kidney disease and the sustainable development goals. Bull. World Health Organ. 96(6):414.

63. Jain S & Chen F (2019) Developmental pathology of congenital kidney and urinary tract anomalies. Clin Kidney J 12(3):382–399.

64. Anand S, Khanam MA, & Finkelstein FO (2014) Global perspective of kidney disease. Nutrition in kidney disease, (Springer), pp 11-23.

65. Murugapoopathy V & Gupta IR (2020) A Primer on Congenital Anomalies of the Kidneys and Urinary Tracts (CAKUT). Clin. J. Am. Soc. Nephrol. 15(5):723–731.

66. Lunt SY & Vander Heiden MG (2011) Aerobic glycolysis: meeting the metabolic requirements of cell proliferation. Annu. Rev. Cell Dev. Biol. 27:441–464.

67. Goyal MS, Hawrylycz M, Miller JA, Snyder AZ, & Raichle ME (2014) Aerobic glycolysis in the human brain is associated with development and neotenous gene expression. Cell Metab. 19(1):49–57.

68. Mizukami Y, et al. (2004) ERK1/2 regulates intracellular ATP levels through α-enolase expression in cardiomyocytes exposed to ischemic hypoxia and reoxygenation. J. Biol. Chem. 279(48):50120–50131.

69. Bhargava P & Schnellmann RG (2017) Mitochondrial energetics in the kidney. Nat Rev Nephrol 13(10):629–646.

70. Emma F, Montini G, Parikh SM, & Salviati L (2016) Mitochondrial dysfunction in inherited renal disease and acute kidney injury. Nat Rev Nephrol 12(5):267–280.

71. Granata S, Dalla Gassa A, Tomei P, Lupo A, & Zaza G (2015) Mitochondria: a new therapeutic target in chronic kidney disease. Nutr. Metab. (Lond.) 12(1):49.

72. Szeto HH (2017) Pharmacologic Approaches to Improve Mitochondrial Function in AKI and CKD. J. Am. Soc. Nephrol. 28(10):2856–2865.

73. Miyazawa H & Aulehla A (2018) Revisiting the role of metabolism during development. Development 145(19):dev131110.

74. Gupta V & Bamezai RN (2010) Human pyruvate kinase M2: a multifunctional protein. Protein Sci. 19(11):2031–2044.

75. Xia J & Wishart DS (2010) MSEA: a web-based tool to identify biologically meaningful patterns in quantitative metabolomic data. Nucleic Acids Res 38(Web Server issue):W71-77.

