## Supplemental figure 1 for "Omics profiling identifies MAPK/ERK pathway as a gatekeeper of nephron progenitor metabolism"

**Supplementary information**

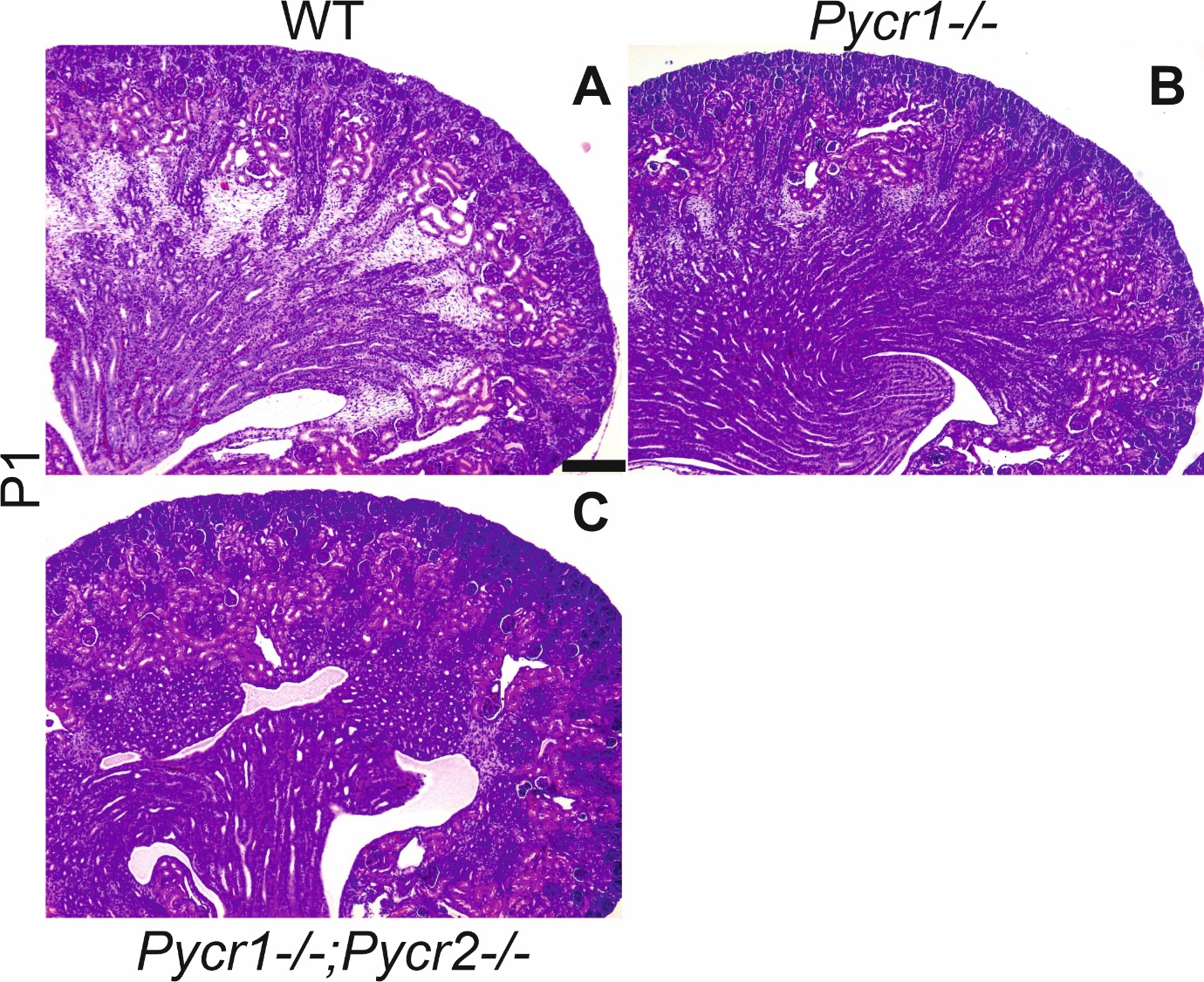

**Supplementary figure 1. Single knockout of *Pycr1* does not affect kidney differentiation.** HE-staining of (**A**) wild type, (**B**) *Pycr1* and (**C**) Pycr1/2 double knockout kidneys at postnatal day 1 (P1). Scale bar: 200µm
